# Informative RNA-base embedding for functional RNA structural alignment and clustering by deep representation learning

**DOI:** 10.1101/2021.08.23.457433

**Authors:** Manato Akiyama, Yasubumi Sakakibara

## Abstract

Effective embedding is being actively conducted by applying deep learning to biomolecular information. Obtaining better embedding enhances the quality of downstream analysis such as DNA sequence motif detection and protein function prediction. In this study, we adopt a pre-training algorithm for the effective embedding of RNA bases to acquire semantically rich representations, and apply it to two fundamental RNA sequence problems: structural alignment and clustering. By using the pre-learning algorithm to embed the four bases of RNA in a position-dependent manner using a large number of RNA sequences from various RNA families, a context-sensitive embedding representation is obtained. As a result, not only base information but also secondary structure and context information of RNA sequences are embedded for each base. We call this “informative base embedding” and use it to achieve accuracy superior to that of existing state-of-the-art methods in RNA structural alignment and RNA family clustering tasks. Furthermore, by performing RNA sequence alignment combining this informative base embedding with a simple Needleman-Wunsch alignment algorithm, we succeed in calculating a structural alignment in a time complexity *O*(*n*^2^) instead of the *O*(*n*^6^) time complexity of Sankoff-style algorithms.

## INTRODUCTION

Effective embedding of DNA sequences, RNA sequences, and amino acid sequences is being actively conducted by using deep representation learning, especially techniques developed in the field of natural language processing (1–3). These studies are based on the idea that nucleotide composition and sequence structure determine the motif and function of a gene sequence, just as the complex grammatical structure of natural language determines the meaning of a sentence. Word embedding techniques for natural language have been applied to nucleotides for DNA sequences. In the dna2vec method (4), word2vec is applied to a DNA sequence to obtain the distributed representation of *k*-mers (DNA sub-sequence of length *k*). Word2vec is one of the effective word embedding techniques (5) that vectorizes the context and meaning of a word using a large amount of text data. Word2vec is based on the hypothesis that words with similar meanings have similar peripheral words. Dna2vec adopts the word2vec technique by defining *k*-mer as a word in the DNA sequence, however dna2vec is hard to be applied to embedding at base-by-base resolution.

Two recently developed state-of-the-art embedding methods are “embeddings from language models” (ELMo) and “bidirectional encoder representations from transformers” (BERT) to generate context-sensitive word-distributed representations (6, 7). In these methods, the same word is assigned different distributed representations that depend on the context. BERT is a pre-training algorithm that obtains word embedding and sentence embedding by performing multiple tasks. The BERT learning algorithm consists of two tasks: a masked language modelling (MLM) task and a next sentence prediction (NSP) task. The MLM task predicts multiple masked tokens (words) in a sentence. The NSP task determines if two statements are consecutive. UniRep (8) and PLUS (9) are representative examples of applying BERT to protein sequence representation; specifically, UniRep obtains the embedding of each amino acid in a protein sequence and uses this embedding to achieve accurate structural and functional predictions of proteins.

In this study, we propose RNABERT for the effective embedding of RNA bases by adopting a pre-learning algorithm BERT to non-coding RNA (ncRNA). We aim to have “informative base embedding” encode the characteristics of each RNA family and structure. To see if the informative base embedding captures the characteristics of RNA family and structure, we apply RNABERT to two basic tasks of RNA sequence analysis: structural alignment and clustering. We evaluate the quality of the informative base embedding by structural alignment of RNA sequences and RNA family clustering.

The first important problem in RNA sequence analysis is the structural alignment of RNA sequences, which calculates the alignment of not only RNA sequences but also their secondary structures. The most influential method for the structural alignment of RNA sequences is the Sankoff algorithm, which simultaneously performs secondary structure prediction and alignment (10). The time complexity of the Sankoff algorithm is *O*(*n*^6^) for the length *n* of input RNA sequences, and accelerating the Sankoff algorithm is an unsolved hard problem (11). While the Sankoff-style algorithms such as LocARNA (12) and Dynalign (13) calculate the alignment considering the secondary structure, a standard sequence-based (non-structural) alignment algorithm such as the Needleman-Wunsch algorithm (14) only determines the correspondence between each base position of two input sequences and its time complexity is *O*(*n*^2^) using the dynamic programming technique. We aim to apply the informative base embedding to determine the position-dependent and secondary structure-dependent match score in calculating alignments so that the structural alignment is obtained using a simple Needleman-Wunsch algorithm instead of a cost-expensive Sankoff-style algorithm.

Since there exist various families of ncRNAs and the high-throughput sequencing continues to generate large number of RNA sequences including novel transcripts, building an appropriate clustering algorithm for ncRNAs is a powerful step towards the discovery of new ncRNA families (15, 16). With the recent increases in deep learning usage, many algorithms for ncRNA classification (supervised clustering) using convolutional neural networks (CNNs) and recurrent neural networks (RNNs) have been proposed (17, 18). These algorithms apply deep learning techniques to one-hot encoding of RNA bases. Most of these algorithms employ supervised learning that uses ncRNA families as labels for training. Since supervised learning requires the data to be labelled, it is not practical when searching for unknown ncRNA families.

For our goals of accurate RNA family clustering and structural alignment of lower computational complexity, we construct an informative base embedding method, RNABERT for RNA sequences that takes into account the context, family, and secondary structure of RNA sequences through two training tasks: MLM and structural alignment learning (SAL). In RNABERT, a pre-training is performed using a large number of unlabelled ncRNA sequences. RNABERT introduces a novel pre-training task, SAL task, in addition to the usual MLM task to more explicitly incorporate RNA secondary structure information into the base embedding for structural alignments. The SAL task employs a pre-training using seed alignments obtained from Rfam database (19) so that bases aligned in the seed structural alignment are expected to have more similar embeddings. We compare the accuracy of structural alignment of RNA sequences between our method and the state-of-the-art methods. Furthermore, by alternately training MLM and SAL tasks, it is expected that RNA-base embedding could adequately capture structural differences among RNA families. We demonstrate that our clustering method is more accurate than existing state-of-the-art methods in clustering of RNA families.

## MATERIAL AND METHODS

### The architecture of RNABERT model

Figure 1 shows the architecture of RNABERT model. RNABERT model consists of three components, token and position embedding, transformer layer, and pre-training tasks. Token embedding randomly generates a 120-dimensional vector representing four RNA bases so that each base is assigned the same vector. Position embedding generates a 120-dimensional vector that represents position information in an RNA sequence. The element-wise sum of token embedding and position embedding for each base in the given RNA sequence is the input vector to the transformer layer. The transformer layer component consists of a stack of 6 transformer layers. Each transformer layer is composed of multi-head self-attention followed by a feedforward neural network. The final output from the transformer layer is an informative base embedding, denoted *Z*. The weights of the transformer layer are trained by alternating two different tasks (MLM and SAL) on top of the output of the transformer layer.

**Figure 1.**
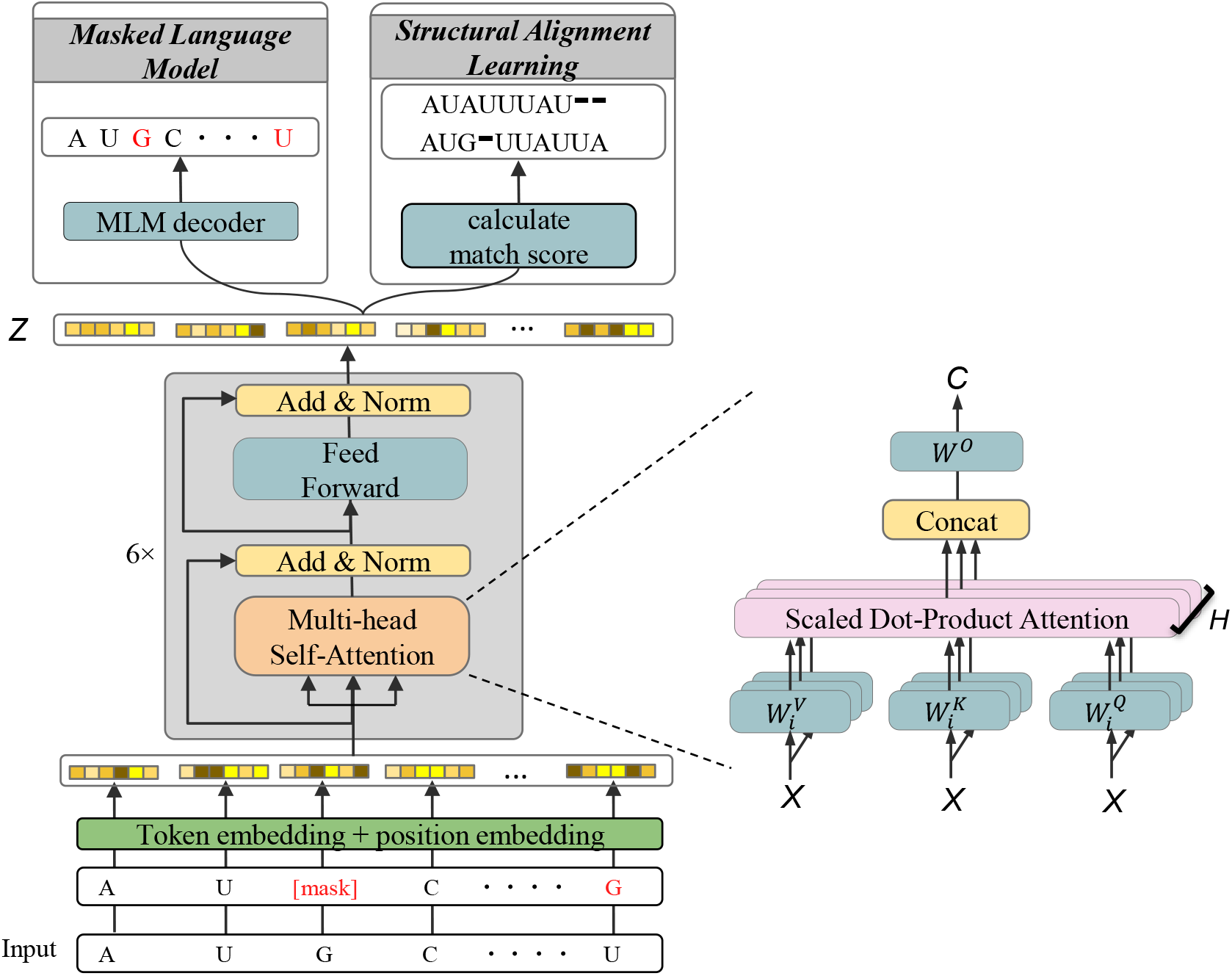
The architecture of RNABERT model. RNABERT model consists of three components, token and position embedding, transformer layer, and pre-training tasks. Token and position embedding randomly generates a 120-dimensional vector representing four RNA bases. The transformer layer component consists of a stack of 6 transformer layers. Each transformer layer is composed of multi-head self-attention followed by a feedforward neural network. The final output from the transformer layer is an informative base embedding, denoted Z. The weights of the transformer layer are trained by alternating two different tasks (MLM and SAL) on top of the output of the transformer layer.

The self-attention mechanism (20) is a central component of the transformer layer. For the transformer layer that takes the output of the previous layer *X* = [*x*_1_, …, *x*_*n*_] as input, the multi-head self-attention with *H* heads compute the output sequence *C* = [*c*_1_, …, *c*_*n*_] with the following formula:

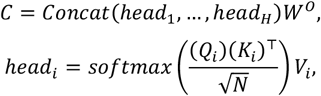

where

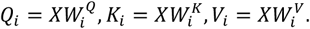

The self-attention mechanism is described as mapping a query and a set of key-value pairs to an output sequence, where the query, key, and value are all matrices: query 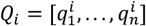, key 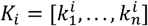 and value 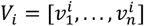. These matrices are the inner products of the learnable weight matrix 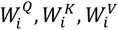 of size *N* × *N*, where *N* is the input and output vector dimension and *N*=120 in our study. In the scaled dot-product attention mechanism, each *head* calculates the next hidden states by computing the attention-weighted sum of the value vector *ν*. An attention coefficient is the output of the softmax function applied to the dot product of the query and key (*Q*_*i*_)(*K*_*i*_)^⊤^ divided by 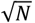. Finally, *H head* ‘s calculated by different set of 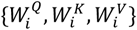 are concatenated and the inner product of this concatenation matrix and *W*^*O*^ yields the output sequence *C*. After the transformer layer process including multi-head attention is performed 6 times, the informative base embedding denoted *Z* is obtained.

### Masked language modelling (MLM)

MLM is a task that masks a part of the input RNA sequence and predicts the masked part using surrounding bases (sub-sequences). The MLM task performs a base embedding so that the masked part can be restored, which enables context-sensitive embedding. First, 15% of bases are randomly selected in a given RNA sequence for training. Next, one of the following three actions is performed on the selected base: 80% of the selected bases are replaced with a token indicating an unspecified base (denoted [mask] in Figure 1), 10% are randomly substituted with one of the other three bases, and the remaining 10% of selected bases are unchanged from their original base. The MLM task trains the model to maximize the probability of correctly predicting the selected 15% RNA bases. In this training model, a classification layer is built on top of the output of the transformer layer. Finally, the output probability of each base is calculated using the softmax function. The cross-entropy function is used as the loss function. Pre-training set for MLM task consists of 762,370 sequences that were generated from those 76,237 human ncRNA sequences obtained from RNAcentral (21) by taking 10 copies of each ncRNA and applying 10 different mask patterns to each.

### Structural alignment learning (SAL)

The SAL task performs a base embedding task to learn the relationship between two RNA sequences. The SAL task is based on RNA structural alignment. RNA structural alignment aligns multiple RNA sequences by inserting gaps between bases so that the conserved secondary structures are aligned in the same column. The SAL task aims to obtain closer embeddings for bases in the same column of reference alignment, and obtain secondary structure embeddings by training based on RNA structural alignment. Rfam seed alignment for each family is downloaded from the Rfam (19) as reference structural alignment for SAL task. To define the loss function in the SAL task, we introduce the *Ω* matrix, which is defined for a pairwise alignment of two RNA sequences and intended to be used as a score matrix when calculating the pairwise alignment. Let *Z* = [*z*_1_, …, *z*_*n*_] and 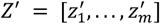 denote the embedded representations output from the transformer layer for the input of two RNA sequences of length *n* and *m*. Each element *ω*_*ij*_ in the *Ω* matrix is defined to be the normalized inner product between *z*_*i*_ and 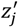.

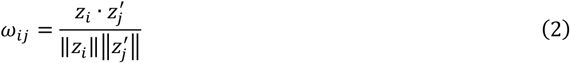

The loss function in the SAL task is defined to increase *ω*_*ij*_ at the match position in the reference alignment so that a sequence alignment algorithm such as the Needleman-Wunsch algorithm produces the reference alignment.

A simple way to implement this loss function in the SAL task is to apply a binary classification learning with respect to *ω*_*ij*_. That is, *ω*_*ij*_ in the aligned position is trained to 1 and *ω*_*ij*_ in unaligned positions is trained to 0. However, this causes strong overfitting. To alleviate this problem, we apply a machine learning method, called “structured support vector machine” (22, 23) to the pre-training in the SAL task. Let an alignment between a pair of RNA sequences *x* = *x*_1_, …, *x*_*n*_ and 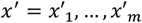 be represented by a series of match (aligned) positions (*i, j*) and gap insertion positions (*i*, −) or (−, *j*) where 1 ≤ *i* ≤ *n*, 1 ≤ *j* ≤ *m*. For a given training dataset *D* consisting of triplets (*x, x*′, *y*) where *x* and *x*′ are a pair of RNA sequences and *y* is the corresponding reference alignment between *x* and *x*′, we aim to find a set of parameters *w* that minimize the following loss function *L*:

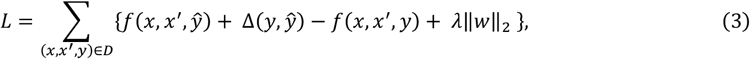

where *f* is the function that returns the score of the alignment *y* between *x* and *x*′. The alignment score is calculated as the sum of the *ω*_*ij*_ value at the match positions (*i, j*) and the gap score at the gap insertion positions (*i*, −) or (−, *j*). *ŷ* is the predicted alignment path calculated by the Needleman–Wunsch algorithm to maximize the sum of the alignment score and the margin term *f*(*x, x*′, *ŷ*) + Δ(*y, ŷ*). The margin term Δ(*y, ŷ*) defines the difference between the reference alignment and the predicted alignment as follows:

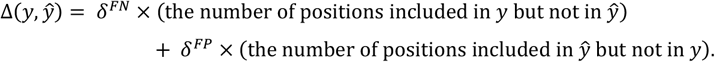

Here, *δ*^*FN*^ and *δ*^*FP*^ are hyperparameters that control the trade-off between sensitivity and specificity for learning parameters. By default, we used *δ*^*FN*^ = 0.05 and *δ*^*FP*^ = 0.1. Decreasing the loss function *L* induces the predicted alignment to be close to the reference alignment.

### RNA structural alignment

A pairwise RNA-sequence alignment based on the base embedding is calculated using the Needleman-Wunsch algorithm using the *Ω* matrix as the score matrix, which is trained in the SAL and MLM tasks. The match score in position (*i, j*) is *ω*_*ij*_ in the score matrix *Ω*, and the gap opening score and gap extension score are set to -1 and -0.1, respectively. The Needleman-Wunsch algorithm, a simple sequence alignment algorithm, is expected to generate RNA structure alignments using informative base embeddings trained in the SAL task. Note that the time complexity of Needleman-Wunsch algorithm is *O*(*n*^2^) for the length *n* of the input RNA sequence.

### RNA family clustering

RNA family clustering is performed as the second evaluation test to confirm the quality of informative base embedding. First, a new measure of similarity between two RNA sequences with respect to soft symmetric alignment (24) is defined as follows. Let *Z* = [*z*_1_, …, *z*_*n*_] and 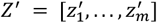 denote the embedded representations output from the transformer layer for the input of a pair of RNA sequences of length *n* and *m*. The similarity *Ŝ* between two RNA sequences is defined to be a weighted sum of the normalized inner product between all *z*_*i*_ and 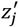 pairs, with the following formula.

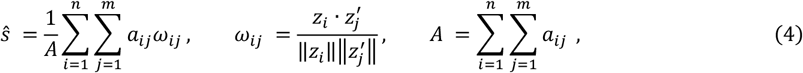

where *a*_*ij*_ is

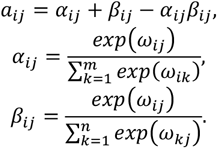

Second, the similarity *Ŝ* is calculated for all pairs of ncRNA sequences to be clustered, and a classification matrix of size *N* × *N* is created, where *N* is the number of RNA sequences in the test dataset. We applied spectral clustering to the rows of the classification matrix by considering each row of the *N*-dimensional vector as a cluster indicator. To confirm the improvement in embedding quality by the SAL task, we compared the clustering accuracy when using only the MLM task and when using the two tasks together.

### Existing methods for RNA structural alignment

There is a family of Sankoff-style algorithms for structural pairwise alignment, which simultaneously solve the sequence alignment problem and the consensus RNA secondary structure prediction problem through dynamic programming. For example, Dynalign and Foldalign (13, 25) use thermodynamic models to find the structure with the lowest free energy common to RNA sequences, while PARTS (26) uses a probabilistic model based on the pseudo-energy obtained from base-pairing probabilities and alignment probabilities to find the most likely structural alignment. While Sankoff-style algorithms yield high alignment accuracy, they are computationally expensive, being *O*(*n*^6^) in time complexity for RNA sequences of length *n*. PMcomp takes base-pairing probability matrices generated using McCaskill’s algorithm as input and incorporates the energy information of each sequence into these matrices to quickly find common secondary structures and alignments (27). Although LocARNA (12) is based on the PMcomp model, the time complexity *O*(*n*^4^) is achieved by simplifying the dynamic programming method utilizing the fact that the base-pairing probability matrix is actually sparse. SPARSE (28) further takes advantage of the sparsity based on the conditional probabilities of bases and base-pairs in the loop region of RNA secondary structure, improving computational time quadratically over LocARNA. RAF (29) achieves the same time complexity as SPARSE by utilizing the sparseness of alignment candidates.

As the baseline for RNA-sequence alignment, a sequence-based alignment algorithm, Needleman-Wunsch algorithm with RIBOSUM score (base substitution) matrix is adopted, which does not take RNA structure information into account. The RIBOSUM 85-60 score matrix (30) is used as a match score between two bases, and the opening gap score and extension gap score are set to -3 and -1, respectively.

### Existing methods for RNA family clustering

The clustering accuracies with state-of-the-art methods GraphClust (15), EnsembleClust (16), and CNNclust (18) were compared. CNNclust is a deep learning-based algorithm that performs supervised learning in which the RNA family class is given as a label. CNNclust can classify RNA families that were not used for training by calculating the similarity score matrix for all pairs of input sequences. We performed experiments with CNNclust using different RNA family groups between training and testing. GraphClust is an unsupervised learning algorithm that does not require the RNA family class as a label and achieves alignment-free clustering with some exceptions. GraphClust employs a graph kernel approach to obtain feature vectors that contain both sequence and secondary structure information. These vectors representing RNA sequences are clustered with a linear time complexity over the number of sequences using a hashing technique. EnsembleClust calculates the similarity between two ncRNAs using expected structural alignment and then applies hierarchical clustering based on the similarity.

### Sequence motif detection using self-attention mechanism

We extracted sequence motifs specific to each RNA family by focusing on the self-attention mechanism. Self-attention is a mechanism used to determine where to focus on the input embedding vectors *X* = [*x*_1_, …, *x*_*n*_] of the input RNA sequence *x* = *x*_1_, …, *x*_*n*_ when generating the output sequence. The attention coefficient sequence *M* = [*m*_1_, …, *m*_*n*_], called attention map, is defined as follows:

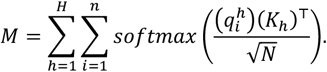

The position *i* with a large *m*_*i*_ value is identified as a motif. Thus, the attention map induces the discovery of the sequence motif, since it indicates a base that is important for training tasks.

### Measures for accuracies of alignment and clustering

Structural alignment accuracy was measured using the sensitivity, positive predictive value, and F1 score, which are calculated as follows. The number of true positives (TP) (or false positives (FP)) is the number of positions (*i, j*) in the predicted alignment that belong (or do not belong) to the reference alignment. The sensitivity of the predicted alignment is TP divided by the number of positions in the reference alignment, and the positive predictive value (PPV) is TP divided by the number of positions in the predicted alignment. The F1 score is the harmonic mean of sensitivity and PPV.

Clustering accuracy was measured with the Rfam family as the true reference class. Three indices, the adjusted Rand index (ARI), homogeneity, and completeness, were used for clustering performance evaluation. The ARI is a measure of how well two types of clustering results match. ARI takes a real number from -1 to 1. If the value of ARI is -1, the two clustering results do not match at all, and if the value is 1, it indicates that they completely match. In this study, ARI shows how close the predicted clustering result is to the true reference class composed of the Rfam family.

The ARI is derived from the Rand index (RI), defined as follows:

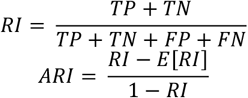

where TP is the number of RNA sequences of the same Rfam family in the same predicted cluster, TN is the number of RNA sequences of a different Rfam family in different predicted clusters, FP is the number of RNA sequences of different Rfam families in the same predicted cluster, and FN is the number of RNA sequences of the same Rfam family in different predicted clusters. Homogeneity is a measure of the proportion of RNA sequences of a single Rfam family that belong to a single predicted cluster. Completeness measures the proportion of RNA sequences of a particular Rfam family that are assigned to the same predicted cluster.

### Datasets

For MLM task pre-training, 76,237 human-derived small ncRNAs of length from 20 to 440 bases from RNAcentral (21) were utilized.

In the training of SAL task, pairwise structural alignment extracted from Rfam seed alignment (19) was used. Ten pairwise alignments were sampled for each seed RNA sequence from 36 RNA families (5.8S rRNA, 5S rRNA, Cobalamin, Entero 5 CRE, Entero CRE, Entero OriR, gcvT, Hammerhead 1, Hammerhead 3, HCV SLIV, HCV SLVII, HepC CRE, Histone3, HIV FE, HIV GSL3, HIV PBS, Intron gpII, IRES HCV, IRES Picorna, K chan RES, Lysine, TAR, Retroviral psi, S box, SECIS, sno 14q I II, SRP bact, SRP euk arch, T-box, THI, tRNA, U1, U2, U6, UnaL2, yybP-ykoY).

For the structural alignment benchmark, we utilized the BRAliBase2.1 k2 database used in the previous study as the gold standard benchmark dataset. Sequence pairs containing unknown bases were eliminated. A total of 8,587 RNA sequence pairs with an average length around 100 bases were used for the benchmark dataset. No alignment overlapped between the training dataset of SAL task and the benchmark test dataset.

To evaluate the clustering accuracy of RNABERT, the test dataset was collected from the BRAliBase2.1 database. The multiple alignment of each ncRNA family provided by the database was treated as a true reference cluster, and each ncRNA sequence in the multiple alignment was treated as a member sequence. All reference clusters with sequence identity less than 70% were selected. The dataset contains the 86 RNA sequences and the 18 RNA families. The RNA sequences used in RNA family clustering test did not overlap with the one used for the training of SAL task.

## RESULTS

### Implementation

RNABERT model was implemented using PyTorch for deep learning. All experiments were run on Linux Red Hat 4.8.5-2 (GPU: tesla v100, CPU: Intel(R) Xeon(R) Gold 6148).

Optuna (31) was used to find the optimal hyperparameters for the MLM task. The list of hyperparameters optimized for the transformer layer are the number of attention heads, the number of transformer layers, feature size, activation function, and training algorithms including Adam, AdaGrad, Momentum SGD. In the MLM task, 5-fold cross-validation was performed, and hyperparameters were determined to maximize accuracy.

### Pretraining of base embedding encodes properties of RNA secondary structure

To investigate whether RNABERT acquired an informative base embedding to encode four RNA bases and secondary structure information, the embedded representations output from the transformer layer for a set of RNA sequences were projected into two-dimensional space using t-SNE (32). In Figure 2, the embedding space adequately represents clusters for four RNA bases (Figure 2, left) and subclusters for characteristic secondary structures (Figure 2, right). Figure 2 shows that RNA base embedding is globally separated by four RNA bases and locally by secondary substructures within each RNA base.

**Figure 2.**
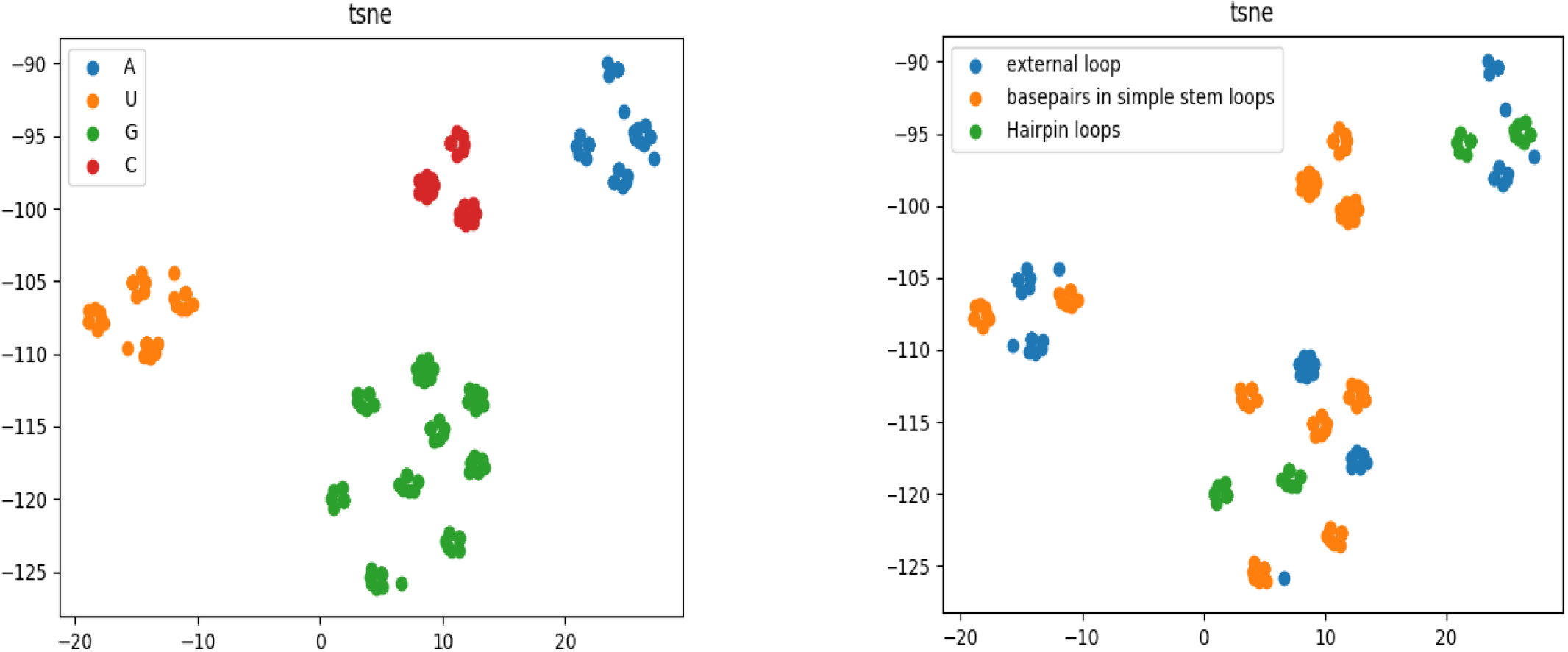
Visualization of RNA base embedding with t-SNE. Shown is a t-SNE projection from a 120-dimensional embedded space into a two-dimensional space. RNA base embeddings were visualized with colours according to the type of RNA-base (top) and the type of characteristic secondary substructure (bottom).

### RNA structural alignment result

Table 1 summarizes the performance evaluation results based on BRAliBase2.1 k2 database for our RNA structural alignment method, denoted RNABERT, and state-of-the-art algorithms for RNA sequence alignment. As shown in Table 1, RNABERT outperformed existing state-of-the-art structural alignment algorithms in all three measures of accuracy.

**Table 1.** RNA structural alignment accuracy and computational time (shown in seconds) of RNABERT, and state-of-the-art algorithms.

|  | Sensitivity | PPV | F1 | Time (sec) |
| --- | --- | --- | --- | --- |
| BERTRNA | **0.881** | **0.947** | **0.913** | 288 |
| LocaRNA | 0.862 | 0.922 | 0.891 | 13,221 |
| SPARSE | 0.848 | 0.931 | 0.888 | 4,216 |
| RAF | 0.865 | 0.938 | 0.900 | 1,423 |
| PARTS | 0.860 | 0.931 | 0.894 | 432,585 |
| Dyalign | 0.706 | 0.914 | 0.797 | 601,226 |
| Sequene-only | 0.632 | 0.796 | 0.705 | **139** |

In terms of computation time, RNABERT was faster than existing state-of-the-art algorithms and was as fast as a sequence-based (non-structural) alignment algorithm with the same computational complexity. The alignment computation of RNABERT consists of three sub-procedures: the first procedure (transformer) obtains the embedding of each base; the second procedure calculates the match score of the two input sequences, and the third procedure calculates the alignment by the Needleman-Wunsch algorithm. The first two procedures can be accelerated by GPU computation. The Needleman-Wunsch algorithm is a simple algorithm that requires *O*(*n*^2^) computation time for two sequences of length n. We achieved high speed computation by implementing the deep learning algorithm using Python and PyTorch, while implementing the Needleman-Wunsch algorithm in the C ++ language.

Figure 3 shows the sensitivity (denoted SEN) and PPV curves calculated for each algorithm for RNA sequence alignment. These values were plotted by sequence identity. As shown in Figure 3, RNABERT yielded very accurate structural alignments and outperformed existing state-of-the-art structural alignment algorithms where sequence identity exceeded 50%. At lower sequence identities, the alignment accuracy of RNABERT was slightly lower than LocARNA and SPARSE, which required larger computation times, and was higher than RAF, which has the fastest computational time among existing structural alignment algorithms. All existing algorithms conduct RNA secondary structure predictions to calculate distances and similarities between RNA sequences. On the other hand, RNABERT does not explicitly use secondary structure predictions, which implies that RNA-base embedding efficiently captures structural information. Especially, for sequences with very low sequence identity, the accuracy of sequence-based alignment tends to decrease, while RNABERT and existing algorithms maintain high accuracy.

**Figure 3.**
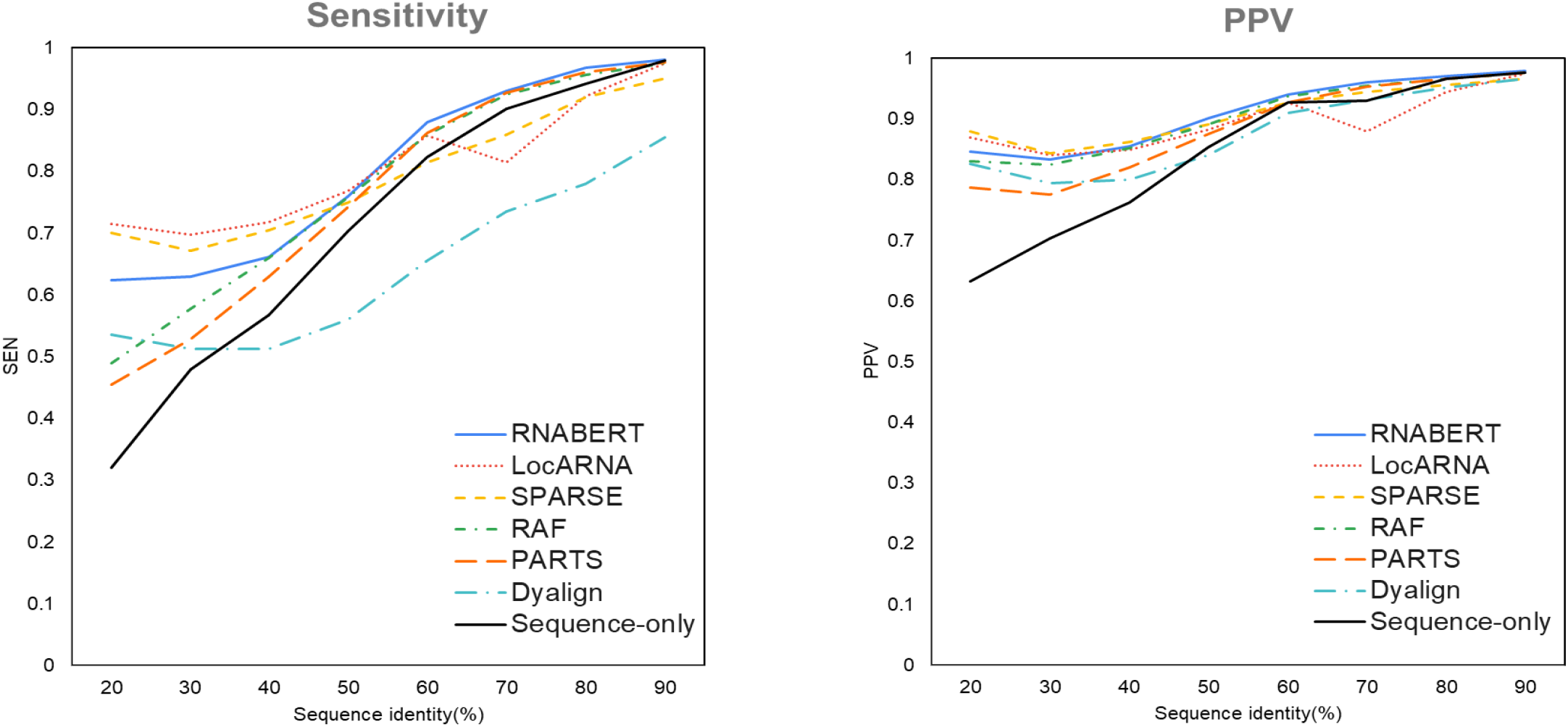
Sensitivity (SEN) and PPV score plots for pairwise RNA structural alignments using RNABERT, LocARNA, SPARSE, RAF, PARTS, Dynalign, and sequence-based (non-structural) alignment with RIBOSUM score matrix (denoted Sequence-only).

### RNA family clustering results

Table 2 shows the ARI, homogeneity and completeness of our RNA clustering method, denoted RNABERT, and state-of-the-art tools for RNA family clustering. RNABERT with MLM and SAL tasks achieved highest ARI and completeness among all state-of-the-art tools. All existing methods use RNA secondary structure prediction to calculate distances and similarities between RNA sequences. This suggests that base embedding by RNABERT, which does not explicitly use secondary structure prediction, efficiently captures structural information. GraphClust and EnsembleClust are based on unsupervised learning and thus do not use the RNA families as labels. The main differences between RNABERT and other methods are the presence of base embedding by pre-learning and the measurement of the similarity between RNA sequences. Since CNNclust is a machine learning-based method that does not use pre-learning unlike RNABERT, this result suggests the effectiveness of base embedding. RNABERT uses soft symmetric alignment as a similarity index between RNA sequences, while GraphClust uses the distance between predicted RNA secondary structures. This result suggests that soft symmetric alignment is an appropriate index of similarity between RNA sequences.

**Table 2.** RNA family clustering accuracy. ARI, homogeneity and completeness of RNABERT, and state-of-the-art tools for RNA family clustering are shown.

|  | ARI | Homogeneity | Completeness |
| --- | --- | --- | --- |
| RNABERT<br>(MLM + SAL) | **0.632** | **0.853** | **0.832** |
| RNABERT<br>(MLM) | 0.465 | 0.722 | 0.609 |
| CNNclust | 0.543 | 0.795 | 0.643 |
| EnsembleClust | 0.596 | 0.667 | 0.664 |
| GraphClust | 0.606 | 0.705 | 0.671 |

### RNA motif

Several well-known sequence motifs in the snoRNA and tRNA families were identified by observing the attention map. Attention maps were extracted from the final transformer layer of RNABERT and indicate the ratios of contribution to the MLM task, and sequence motifs were detected from the attention map. The “UUCGA” sequence motif shown in Figure 4a is typical in the T loop of tRNA (33). The motif is specifically present in TRT-AGT6-1, as displayed in the secondary structure in Figure 4b. The motifs depicted in Figure 4c are the typical motifs “UGAUGA” and “CUGA” present in the snoRNA C/D box (34, 35). The motif is specifically present at SNORD113-7, as displayed in the secondary structure in Figure 4d.

**Figure 4.**
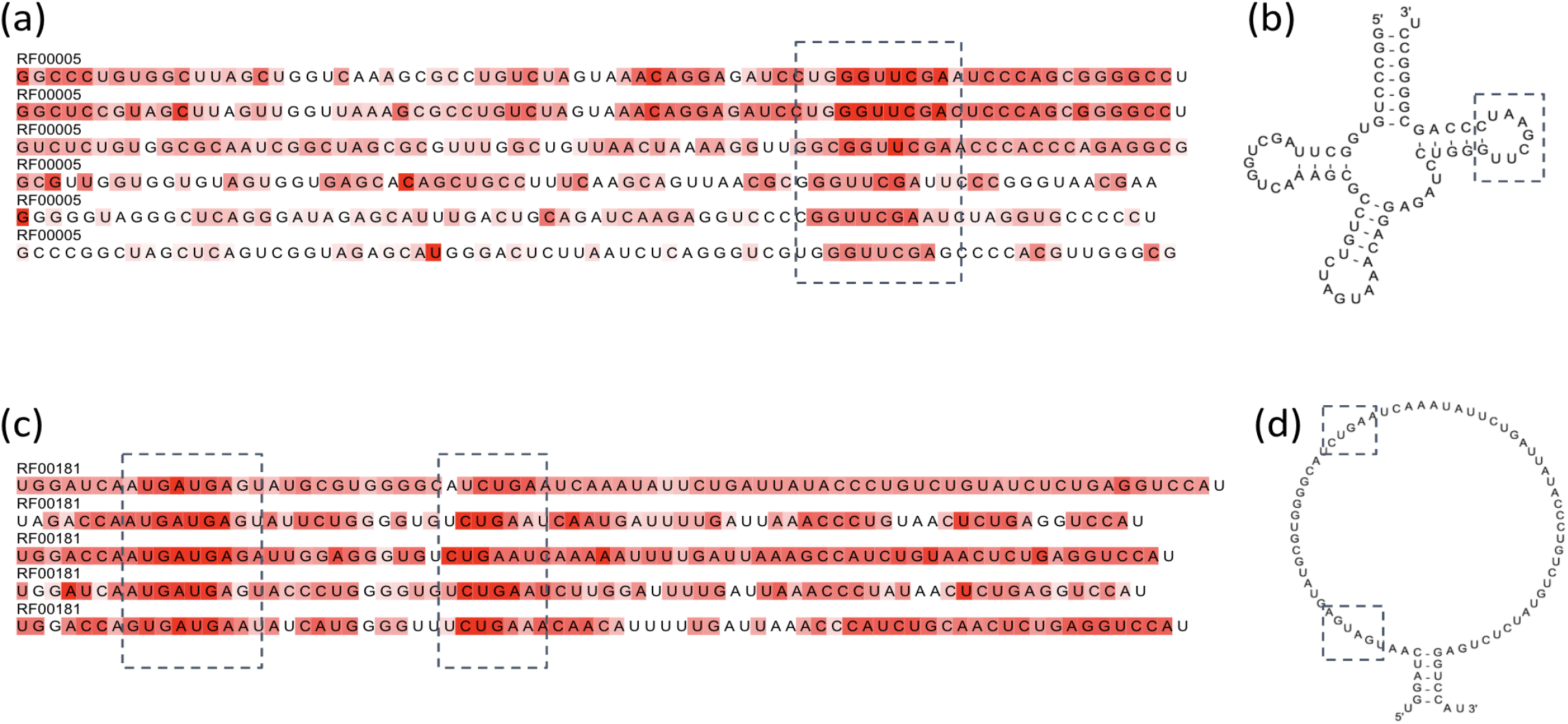
Extracted sequence motifs of the tRNA ((a), (b)) and snoRNA families ((c), (d)). (a) and (c) are visualizations of the attention map at each base. Bases with darker red background have higher attention map values.

## DISCUSSION

We performed three tasks to obtain “informative base embedding”. The MLM task is a fundamental task developed in the original BERT, and the SAL is a novel RNA-sequence specific task introduced in this study. To see if these tasks effectively incorporate RNA secondary structure information into base embedding, we performed two tests, RNA clustering and sequence alignment. Compared to other existing algorithms, RNABERT achieved high accuracy in family clustering. This indicates that RNABERT effectively performed RNA family classification by pre-training with a large number of RNA sequences and has a high ability to cluster unknown RNA families. With the development of high-throughput sequencing, hundreds of thousands of ncRNAs have been detected, but many have not been annotated. In fact, 86% (24,972,896) of the 28,895,596 ncRNAs present in RNAcentral do not have GO annotation. RNABERT can be applied to the detection of these unknown RNA families.

Various methods have been developed for RNA structural alignment, and Sankoff-style algorithms that simultaneously predict optimal alignment and folding are actively studied. Such Sankoff-style algorithms are known to give accurate structural alignment results, especially for RNAs with relatively low sequence similarity, but the algorithms are usually very complex in time and space. Unlike the many structural alignment algorithms based on the Sankoff algorithm, RNABERT does not explicitly consider RNA folding, but boasts a high degree of structural alignment accuracy. This is considered to be evidence that the base embedding encodes the secondary structure information specific to RNAs. Furthermore, while RNABERT achieves the same accuracy as Sankoff-style algorithms, it is much faster because it uses a simple sequence-based alignment algorithm. In fact, the time complexity of the RNABERT algorithm is *O*(*n*^2^) for two sequences of length *n*.

SPARSE (28) addressed an improvement of computational time for Sankoff-style algorithm. SPARSE achieved quadratic time for simultaneous alignment and folding by assuming the sparse structure on RNA secondary structures. On the other hand, RNABERT achieved quadratic computational time by reducing the problem of RNA structural alignment problem into the problem of sequence alignment based on pretraining of base embedding. In fact, the computational time of RNABERT was an order of magnitude faster than that of SPARSE, as revealed in this study.

The base embedding obtained by RNABERT is applicable to various fields for RNA informatics. The most immediate problem is the extension to the multiple structural alignment of RNA sequences. It is expected to accomplish this task by combining existing sequence-based multiple alignment algorithms such as MUSCLE (36) and MAFFT (37) with the score matrix *Ω* and the informative base embedding. Another area most likely to improve with the application of RNABERT is RNA secondary structure prediction. Since the embedding of the base contains information on the secondary structure, it can be expected to contribute to the prediction of the RNA secondary structure (38). Similarly, base embedding can be applied to RNA interactome (RNA-protein interaction, RNA-RNA interaction) in which the RNA secondary structure acts on the interaction between molecules. This study has not addressed RNA modification (e.g., m6A, m1A), but it may be helpful to use this information for more precise modelling of base embedding.

## AVAILABILITY

The codes, datasets and pre-trained RNABERT models are available at https://github.com/mana438/RNABERT.git.

## LIST OF ABBREVIATIONS

BERT: Bidirectional Encoder Representations from Transformers
MLM: Masked Language Modelling
ncRNA: non-coding RNA
CNN: Convolutional Neural Network
SAL: Structural Alignment Learning
ARI: Adjusted Rand Index

## AUTHORS’ CONTRIBUTIONS

MA; implemented the software, analysed data, compared with the existing methods, and co-wrote the paper. YS; designed and supervised the research, analysed data, and co-wrote the paper. All authors read and approved the final manuscript.

